# Resolving an underrepresented circulating tumor cell population in lung cancer enabled by Hexokinase 2 analysis

**DOI:** 10.1101/2020.04.27.064345

**Authors:** Liu Yang, Xiaowei Yan, Jie Chen, Qiong Zhan, Yingqi Hua, Shili Xu, Yu Dong, Ziming Li, Zhuo Wang, Dongqing Zuo, Min Xue, Yin Tang, Harvey R. Herschman, Shun Lu, Qihui Shi, Wei Wei

**Author notes:** These authors contributed equally to this work (Liu Yang, Xiaowei Yan, Jie Chen, Qiong Zhan, Yingqi Hua). Correspondence and requests for materials should be addressed to S. L. or Q. H. S. or W. W.

## Abstract

Unlike other epithelial cancer types, circulating tumor cells (CTCs) are less frequently detected in the peripheral blood of non-small cell lung cancer (NSCLC) patients using epithelial marker-based detection approaches, despite the aggressive nature of NSCLC. Here we demonstrate hexokinase-2 (HK2) as a metabolic function-associated marker for detection of CTCs, with significantly improved detection rates and high specificity, in 33 NSCLC patients. Use of the HK2 marker identified underrepresented cytokeratin-negative (HK2^high^/CK^neg^) CTCs present in many blood samples but rarely detected in pleural effusions or cerebrospinal fluids of NSCLC patients. HK2^high^/CK^neg^ CTCs exhibited smaller sizes but consistent copy number variation profiles compared to CK^pos^ CTCs. Surprisingly, CK expression levels were found to be independent of CTC epithelial-to-mesenchymal transition (EMT) status as measured by single-cell transcriptome profiling, challenging the long-standing association between CK expression and EMT. Our approach improves sensitivity of CTC detection in NSCLC and can potentially resolve a more complete spectrum of CTCs, regardless of their CK expression levels or epithelial traits.

## Introduction

Circulating tumor cells (CTCs) are known to spread from primary sites through blood circulation and other body fluids to seed distant metastases, the main cause of cancer-related deaths (*1, 2*). The presence of CTCs correlates with increased metastatic propensity and burden. Consequently CTCs are widely considered as one of the most promising biomarkers for hematogenous metastases (*2*). Due to the extreme rarity of CTCs and the epithelial nature of many cancers, the epithelial cell adhesion molecule (EpCAM) and different members of cytokeratin (CK) family are currently employed for candidate CTC enrichment and identification, prior to subsequent characterization (*3–5*). The EpCAM/CK-based detection strategy has been adopted by the most commonly used, FDA-cleared CellSearch system for CTC enumeration, although this procedure has intrinsic limitations in detecting CTCs from non-epithelial malignancies (*4*).

Although the CellSearch system provides reliable performance in breast, prostate, and colorectal cancers (*3, 4*), CTCs are much less frequently observed in peripheral blood of non-small cell lung cancer (NSCLC) patients when compared to other epithelial cancers using EpCAM/CK-based methods, despite the highly aggressive nature of NSCLC (*5–8*). Previous studies reported that less than 5 CTCs were detected in 7.5 mL blood in a majority (~85%) of stage IV NSCLC patients, and essentially no CTCs were detected in the blood from early stage patients with localized NSCLC disease (*6, 8*). However, a significant portion of these early stage patients with resectable tumors and few or no detectable CTCs develop metastatic relapse. These data suggest that significant populations of NSCLC CTCs that express low or no EpCAM or CK may exist, but escape detection by these epithelial markers. While the introduction of mesenchymal markers such as N-Cadherin or Vimentin might help increase the coverage, these CK^neg^ (or EpCAM^neg^) CTCs may not have high levels of mesenchymal marker expression. Moreover, the presence of reactive mesenchymal stromal cells, tumor-derived circulating endothelial cells, and hematopoietic cells of mesenchymal origin compromise the specificity of these mesenchymal markers (*3, 9–11*). Ideally, to achieve full-spectrum for CTC detection in NSCLC, a marker that exploits a common feature of all cancer cells is desired.

Our approach to identifying and validating a new CTC marker arises from a characteristic that is associated with an aberrant function present in cancers of many different origins. A key hallmark of many cancers is the capacity to metabolize glucose at an elevated rate. This capacity has been exploited clinically by positron emission tomography (PET) for cancer diagnosis (*12*). The first enzymatic step of glycolysis, critical to the elevated glucose metabolism observed in most cancer cells, is the phosphorylation of glucose catalyzed by hexokinase (HK) (*13*). Four HK enzymes, HK1, HK2, HK3 and HK4 (glucokinase), have been identified in mammals. Glucokinase is expressed primarily in the liver. HK1 is expressed in the cells of many normal tissues. While HK2 is expressed in embryonic tissues and in a few adult tissues (e.g. muscle and adipose), most adult cells express little or undetectable HK2 at physiological conditions. In contrast, HK2 is expressed at substantial levels in a wide range of cancers, including cancers of both epithelial and non-epithelial origins (*13–16*). The high level of HK2 expression and activity in glycolytic tumor cells, revealed in PET imaging through the increased conversion of ^18^F-FDG to ^18^F-FDG-6P, has been associated with poor overall survival in cancer patients (*17*). In addition, the association of HK2 with mitochondria suppresses the death of cancer cells, thus increasing their possibility for metastasis (*13–15*). Given its selective expression in cancer cells and its restricted expression in normal adult tissues, HK2 is a potential marker to discriminate CTCs with elevated glycolysis, independent of their expression levels of commonly used epithelial or mesenchymal markers, from other cell types in liquid biopsies.

Here we investigate the utility of HK2 as a glycolytic activity-associated marker to identify CTCs in liquid biopsies from lung adenocarcinoma (LUAD) patients. The HK2 marker, in combination with CK and CD45, allows immunologic detection of a greater spectrum of CTCs in LUAD patient blood, at a much higher detection rate than can be identified by traditional CellSearch-like systems. In addition to the CK^pos^ CTCs, HK2 analysis can identify an underrepresented HK2^high^/CK^neg^ CTC population that cannot be effectively detected by traditional assays, Analyzing three common types of liquid biopsy samples of LUAD patients using the HK2 marker combination, we found enrichment of HK2^high^/CK^neg^ CTCs in peripheral blood, but not in malignant pleural effusions (MPE) or cerebrospinal fluid (CSF) of LUAD patients. Single-cell sequencing was used to confirm the malignancy of these CTCs by characterizing genome-wide copy number variation (CNV) to validate the specificity of our CTC detection approach. Physical and molecular features were compared at the single-cell level between CK^pos^ and CK^neg^ CTC populations. Our results challenge a long-standing notion that there is a negative correlation between CK expression and epithelial-mesenchymal transition (EMT), providing insights into this underrepresented HK2^high^/CK^neg^ CTC population in lung cancer patients. More generally, the HK2-based approach is likely to be useful in identifying CTCs from patients with a wide variety of cancers.

## Results

### Principle of HK2-based CTC identification

In contrast to normal tissues that primarily use HK1 for glycolysis, many tumor cells express elevated levels of HK2 (*13–16*). High levels of HK2 expression are observed in a wide range of cancer cell lines derived from tissues of different origin (Fig. 1A). HK2 expression, generally believed to be a driver of increased glycolysis in tumors (*13*), is evaluated here to detect CTCs from three types of liquid biopsy samples from lung cancer patients.

**Figure 1.**
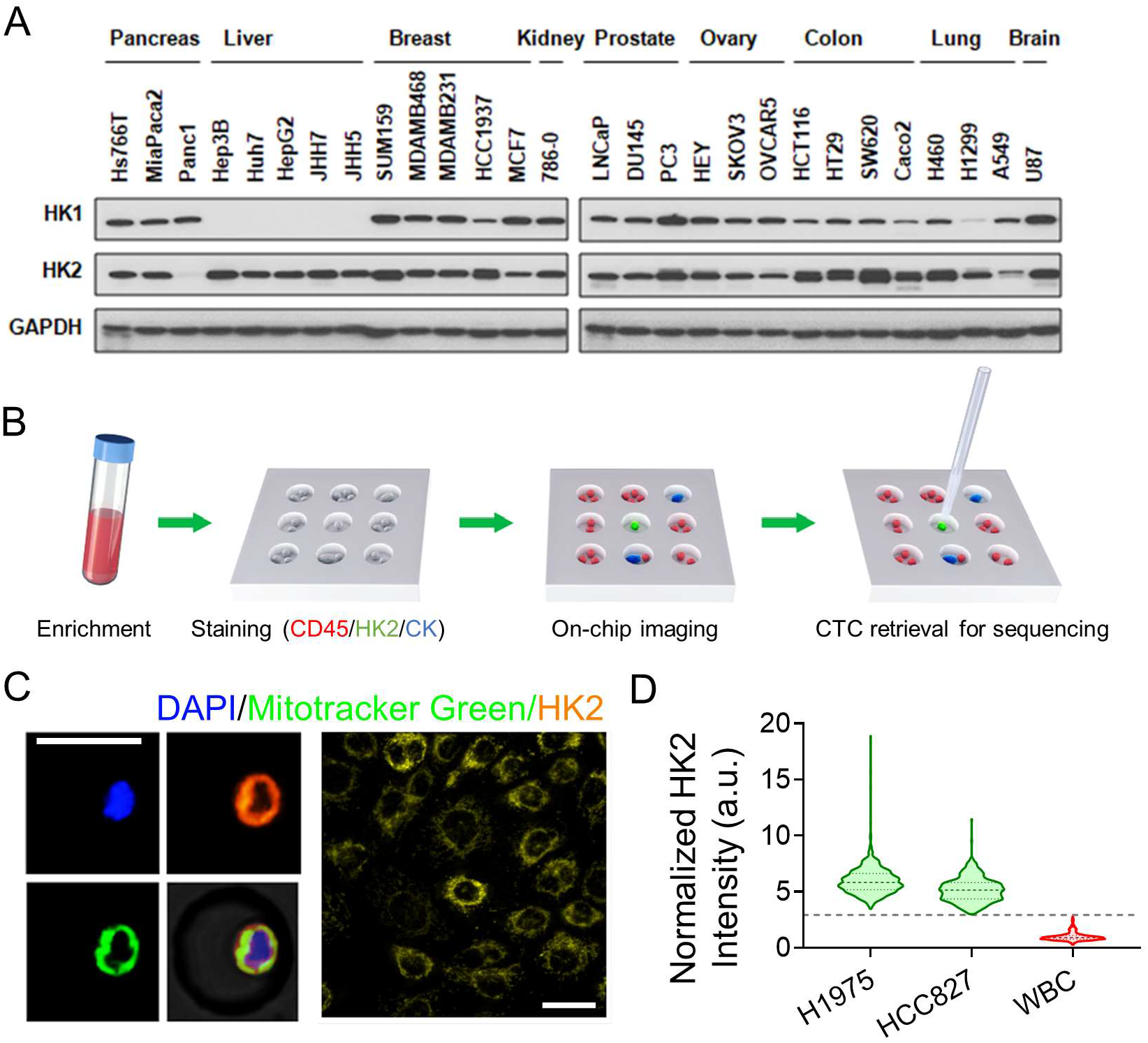
Overall strategy and technical platform validation of HK2-based on-chip image cytometry assay. (A) Immunoblotting results showing HK2 overexpression in a wide range of cancer cell lines. (B) The work flow of CTC enrichment, identification and single-cell manipulation using on-chip image cytometry assay. A PDMS microwell chip was used with each well 30 μm in diameter and 20 μm in height. (C) Left, co-localization of HK2 and MitoTracker Green in H1975 cells. Right, fluorescence image of HK2-stainied H1975 cells at 40× magnification. HK2 staining exhibits a fragmented morphology, with many spheroid-shaped staining, or a reticulated morphology, very similar to the morphology of mitochondria. Scale bar: 30 μm. (D) Violin plots of HK2 intensity of H1975 and HCC827 cells that were normalized to the mean of white blood cells (N=591, 433, and 1027, respectively). The dash and dot lines of each violin plot denote the median as well as first and third quartiles, respectively. The grey dash line indicates the average HK2 intensity of leukocytes plus 5 times the standard deviation.

The HK2-based CTC identification assay is performed on a single-cell on-chip image cytometry platform (Fig. 1B). Briefly, after removal of red blood cells and platelets and depletion of CD45^pos^ cells using a RosetteSep™ CTC Enrichment Cocktail followed by density gradient centrifugation, the enriched cells are recovered and applied into a PDMS microwell chip with thousands of addressable microwells for on-chip cell fixation, permeabilization, and immunostaining of HK2, pan-CK (CK7/8), CD45, and DAPI, followed by washing and imaging. To avoid cell loss during on-chip operations, a porous membrane is used to seal the chip after cell loading, and prior to immunostaining. The chip is imaged by a high-content fluorescent imaging system. A computational algorithm analyzes the images at single-cell resolution, identifies putative CD45^neg^/HK2^high^/CK^pos/neg^ CTCs and CD45^neg^/HK2^low^/CK^pos^ CTCs based on the calculated fluorescence cut-offs, and reports the corresponding microwell addresses. While the CD45^neg^/HK2^low^/CK^neg^ cell population may contain some CTCs with potentially low glycolytic activities, there also exist in this population confounding cell types that include CD45^neg^ immune cells (*e.g.* plasma cells), circulating endothelial cells, mesothelial cells (in MPE), and dying CTCs (fig. S1). Consequently they are excluded from our analysis. All the putative CTCs are then retrieved individually by a motorized micromanipulator, based on recorded microwell addresses, for single-cell sequencing to determine the malignancy of each cell (Fig. 1B).

As a proof-of-concept demonstration, we spiked tumor cells from two representative LUAD cell lines known to express substantial levels of HK2 into 5 mL of healthy donor blood, respectively, then followed by CTC enrichment and the on-chip immunostaining protocol described above. Consistently, we found that HK2 staining co-localized with mitochondrial staining (Fig. 1C and fig. S2) (*13*). The HK2 fluorescence signals of the spiked H1975 and HCC827 cells are approximately 5 fold greater than those of leukocytes (Fig. 1D). A cutoff line defined by the average HK2 signal plus 5 times the standard deviation of CD45^pos^ leukocytes can segregate all leukocytes from spiked cancer cells (Fig. 1D). Therefore, in subsequent analyses of biological fluids from LUAD patients, we designated levels greater or smaller than this cutoff line to be HK2^high^ or HK2^low^. Additionally, leukocytes are normally deemed CK7/8^neg^ (*18*); consequently, their average background signal of CK plus 5 times the standard deviation can cover the signal range of almost all leukocytes and was set as the cut-off for gating CK^neg^ from CK^pos^ cells (Fig. 2A).

**Figure 2.**
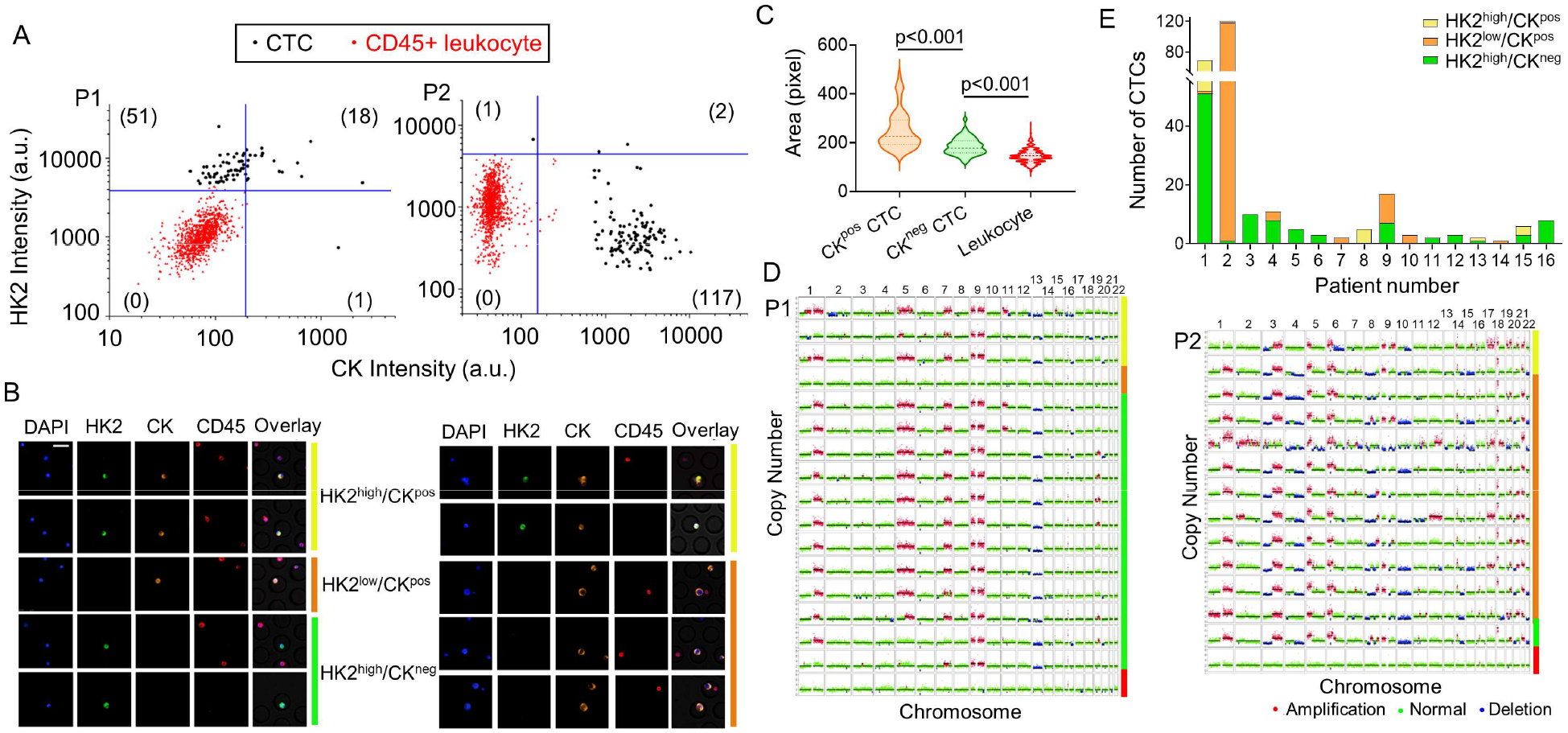
Identification and characterization of CTCs in the blood samples of LUAD patients. (A) Scatter plots report HK2 and CK immunostaining fluorescence intensities of CTCs and CD45^pos^ leukocytes in the blood samples from P1 and P2. HK2^high^ and CK^pos^ cells are gated out by five standard deviations above the mean of HK2 or CK levels of CD45^pos^ leukocytes. Numbers of HK2^high^/CK^neg^/CD45^neg^, HK2^low^/CK^pos^/CD45^neg^, and HK2^high^/CK^pos^/CD45^neg^ CTC subsets are displayed in the figure. (B) Representative fluorescence images of CTCs from P1 (left) and P2 (right). Images for HK2^high^/CK^pos^, HK2^low^/CK^pos^, and HK2^high^/CK^neg^ CTCs are color-coded by yellow, orange, and green bars, to the right, respectively. (C) Comparison of cell sizes among CK^pos^ CTCs, CK^neg^ CTCs, and leukocytes from P1 and P2 (N=138, 52 and 2045, respectively). The dash and dot lines of each violin plot denote the median and first and third quartiles, respectively. (D) Single-cell CNV profiles across the autosomes of randomly selected CTCs and leukocytes from P1 and P2, highlighting their genome-profile similarity independent of CK expression. The CTC subtypes are color-coded to the right in the same way as panel B, and leukocytes are color-coded by red bars. (E) CTC counts and subtypes in a total of 16 LUAD patients with positive CTC counts in this study.

### HK2^high^/CK^neg^ CTCs as a prevalent subtype in half of peripheral blood samples of LUAD patients

Having confirmed that HK2 can detect different LUAD cell lines spiked in the peripheral blood samples, we applied this method to blood samples from LUAD patients. Putative CTCs were first enriched and then detected through a combination of HK2, pan-CK (CK7/8), DAPI, and CD45 staining, followed by validation with single-cell sequencing of genome-wide copy number variation (CNV) profiles. Candidate CTCs were identified based on the marker criteria discussed above. Two typical CTC subtypes based on the CK expression levels were identified in the blood samples. For Patient 1 (P1), a total of 70 putative CTCs were identified in 5 mL of blood, including 51 HK2^high^/CK^neg^/CD45^neg^ cells, 18 HK2^high^/CK^pos^/CD45^neg^ cells and one HK2^low^/CK^pos^/CD45^neg^ cell (Fig. 2, A and B, table S1).These putative CTCs were all DAPI positive and ~73% of them exhibited a CK^neg^ subtype with high HK2 staining. In contrast, for Patient 2 (P2), a total of 120 putative CTCs were identified in 5 mL of blood, including 117 HK2^low^/CK^pos^/CD45^neg^ cells, two HK2^high^/CK^pos^/CD45^neg^ cells and one HK2^high^/CK^neg^/CD45^neg^ cell (Fig. 2, A and B, table S1). Around 99% of detected CTCs exhibited a CK^pos^ subtype and 97.5% showed low HK2 staining. Remarkably, sizes of CK^pos^ CTCs were statistically larger than those of CK^neg^ CTCs in P1 and P2 blood samples (Fig. 2C).

Single-cell CNV analysis allowed us to survey the entire genomic landscape and determine the malignancy of the cell (i.e. the cancer cell genotype) based on chromosomal gains and losses at the single-cell level. The CNV profiles of 16 randomly selected CTCs of P1 displayed largely reproducible copy number alterations across the genome, distinct from leukocytes (Fig. 2D). The copy number gain in chromosome 9 and loss in chromosome 13, in particular, were recurrent across nearly all the putative CTCs sequenced. Such reproducible global CNV profiles were shared in 100% of sequenced HK2^high^ cells, regardless of their CK levels, confirming the tumor origin of these cells. Likewise, all sequenced putative CTCs from P2 also displayed reproducible gain and loss in CNV patterns, independent of CK or HK2 levels (Fig. 2D). The results from both patients validate the high specificity of this marker combination (i.e. CD45^neg^/HK2^high^/CK^pos/neg^ and CD45^neg^/HK2^low^/CK^pos^) in detecting CTCs in LUAD patient blood samples.

We used these marker combinations to interrogate peripheral blood samples from a cohort of 24 treatment naïve LUAD patients with advanced stages (table S1). CTCs were detected in ~67% of patients based on our markers. This percentage is substantially higher than the 20%-39% CTC positivity rates using EpCAM/CK-based approaches reported in previous studies (*19*). A CTC number ≥ 5 in 5 mL of blood was detected in ~37% patients. In contrast, a CTC number ≥ 5 in 7.5 mL of blood that was detected in only 9% of patients in a previous study (*6*). We conclude that the use of the HK2/CK/CD45 marker combination can achieve a much higher CTC detection sensitivity than traditional EpCAM/CK-based approaches. The malignancy of the putative CTCs was confirmed by single-cell CNV analysis of randomly selected CTCs, validating the assay specificity with single-cell precision. Interestingly, while the CNV profiles of CTCs from the same patient displayed a high consistency (Fig. 2D), CTCs from different patients exhibited diverse CNV patterns, highlighting the interpatient heterogeneity of these CTCs (fig. S3).

For patients with positive CTC counts, HK2^high^/CK^neg^ CTCs were detected in 75% of these patients and were the prevalent phenotype (> 50% of total CTCs in a patient) in 50% of them (Fig. 2E). Around ~37% of the patients with positive CTCs counts had only HK2^high^/CK^neg^ CTCs detected in their blood samples. These cells were not accessible to EpCAM/CK-based detection approaches. Consequently, our increased sensitivity for CTC detection in LUAD patients is mainly attributed to the introduction of the glycolytic activity-associated HK2 marker. These results prompted us to investigate whether other types of liquid biopsies from LUAD patients, such as MPE or CSF, contain a significant portion of CK^neg^ CTCs.

### HK2^high^/CK^neg^ CTCs as a minority subtype in MPE and CSF samples from LUAD patients

HK2^high^/CK^neg^ CTCs were found not only in peripheral blood but also other types of liquid biopsies, such as MPE and CSF, of LUAD patients. For lung cancer patients, tumor cells present in pleural effusion denotes an advanced stage of disease with metastasis (M1a staging) (*20*). We did not assess HK2^low^/CK^pos^/CD45^neg^ cells because, in pleural effusions, there are significant numbers of detached mesothelial cells that also express CK7/8 but have low HK2 levels (in contrast to blood samples). In 10 mL of the pleural effusion sample from P26, we identified a total of 367 putative CTCs based on the HK2^high^/CD45^neg^/DAPI^pos^ definition. These cells were classified into two categories that include 355 HK2^high^/CK^pos^/CD45^neg^ CTCs and 12 HK2^high^/CK^neg^/CD45^neg^ CTCs, based on their CK levels (Fig. 4A and table S2). Single-cell genome analysis demonstrated reproducible CNV profiles across all the CTCs sequenced, regardless of their CK expression levels (Fig. 4B). Similar CNV profile consistency was also observed among CTCs in the MPE sample of P25, while to a lesser degree (fig. S4). These results confirm HK2 as a reliable marker of CTCs independent of CK expression, which is especially useful to identify those CK^neg^ CTC populations. In contrast to blood samples, across the 6 MPE samples analyzed from LUAD patients, we found that HK2^high^/CK^neg^/CD45^neg^ CTCs constituted a minority population in the pleural effusions, ranging from 0.2% to 20%; HK2^high^/CK^pos^/CD45^neg^ CTCs are the majority population (Fig. 3C and table S2).

**Figure 3.**
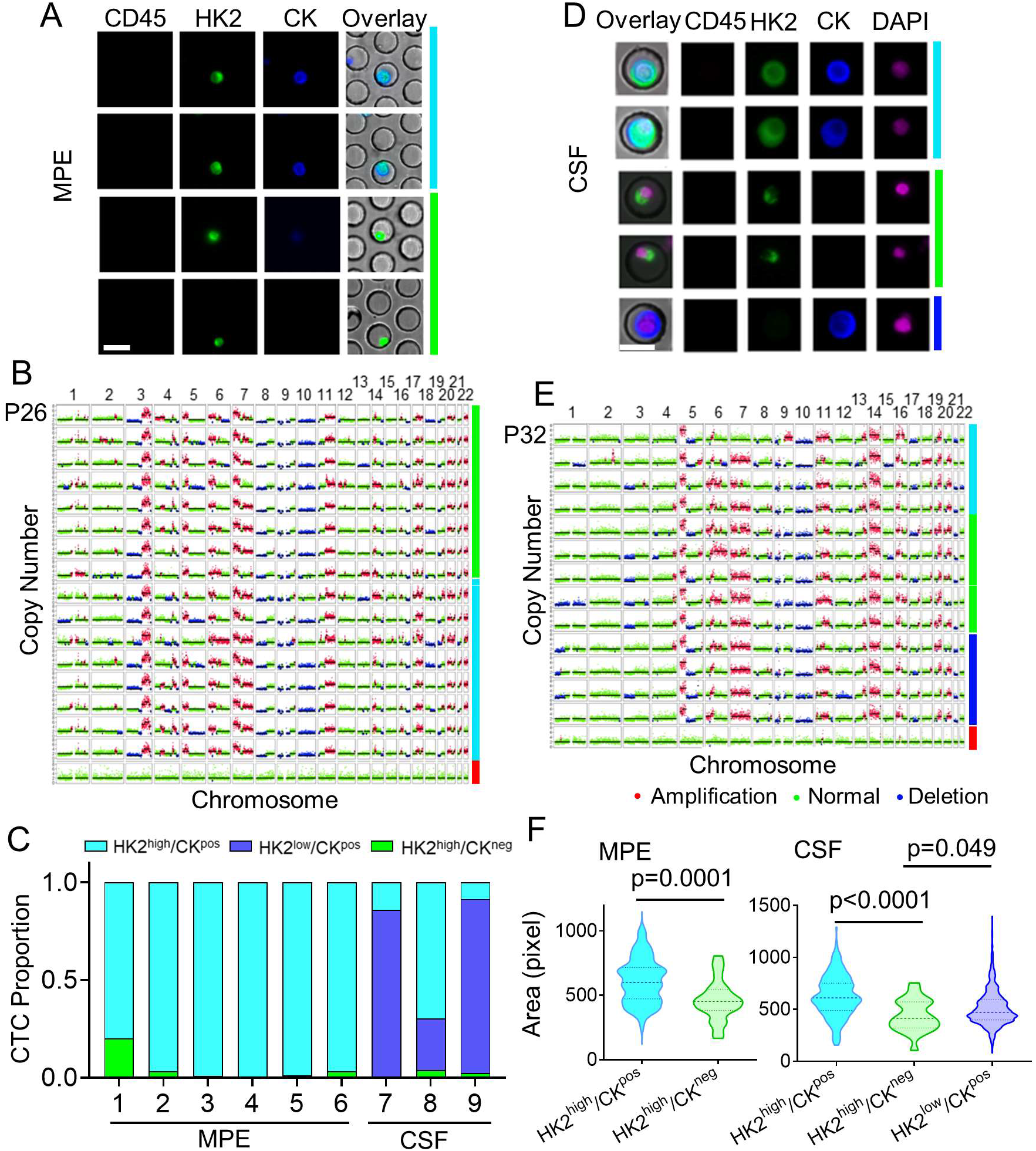
Identification and characterization of CTCs in the MPE and CSF samples of LUAD patients. (A) Representative fluorescence images of CTCs from an MPE sample of P26. Images for HK2^high^/CK^pos^/CD45^neg^ and HK2^high^/CK^neg^/CD45^neg^ CTCs are color-coded by cyan and green bars to the right, respectively. (B) Single-cell CNV profiles of randomly selected CTCs and leukocytes from the MPE sample of P26. The CTC subtypes are color-coded to the right in the same way as panel A and leukocytes are color coded by red bars. (C) CTC subtype proportions across all the MPE and CSF samples. (D) Representative fluorescence images of CTCs from a CSF sample of P31. Images for HK2^high^/CK^pos^/CD45^neg^, HK2^high^/CK^neg^/CD45^neg^, and HK2^low^/CK^pos^/CD45^neg^ CTCs are color-coded by cyan, green, and blue bars to the right, respectively. (E) Single-cell CNV profiles of randomly selected CTCs and leukocytes from the CSF sample of P31. The CTC subtypes are color-coded to the right in the same way as panel D and leukocytes are color coded by red bars. (F) Comparison of cell sizes between HK2^high^/CK^pos^ and HK2^high^/CK^neg^ CTCs in the MPE sample (left, N=100 and 25, respectively) and HK2^high^/CK^pos^, HK2^high^/CK^neg^, and HK2^low^/CK^pos^ CTCs in the CSF sample (right, N=258, 22 and 1498, respectively). The dash and dot lines of each violin plot denote the median and first and third quartiles, respectively.

**Figure 4.**
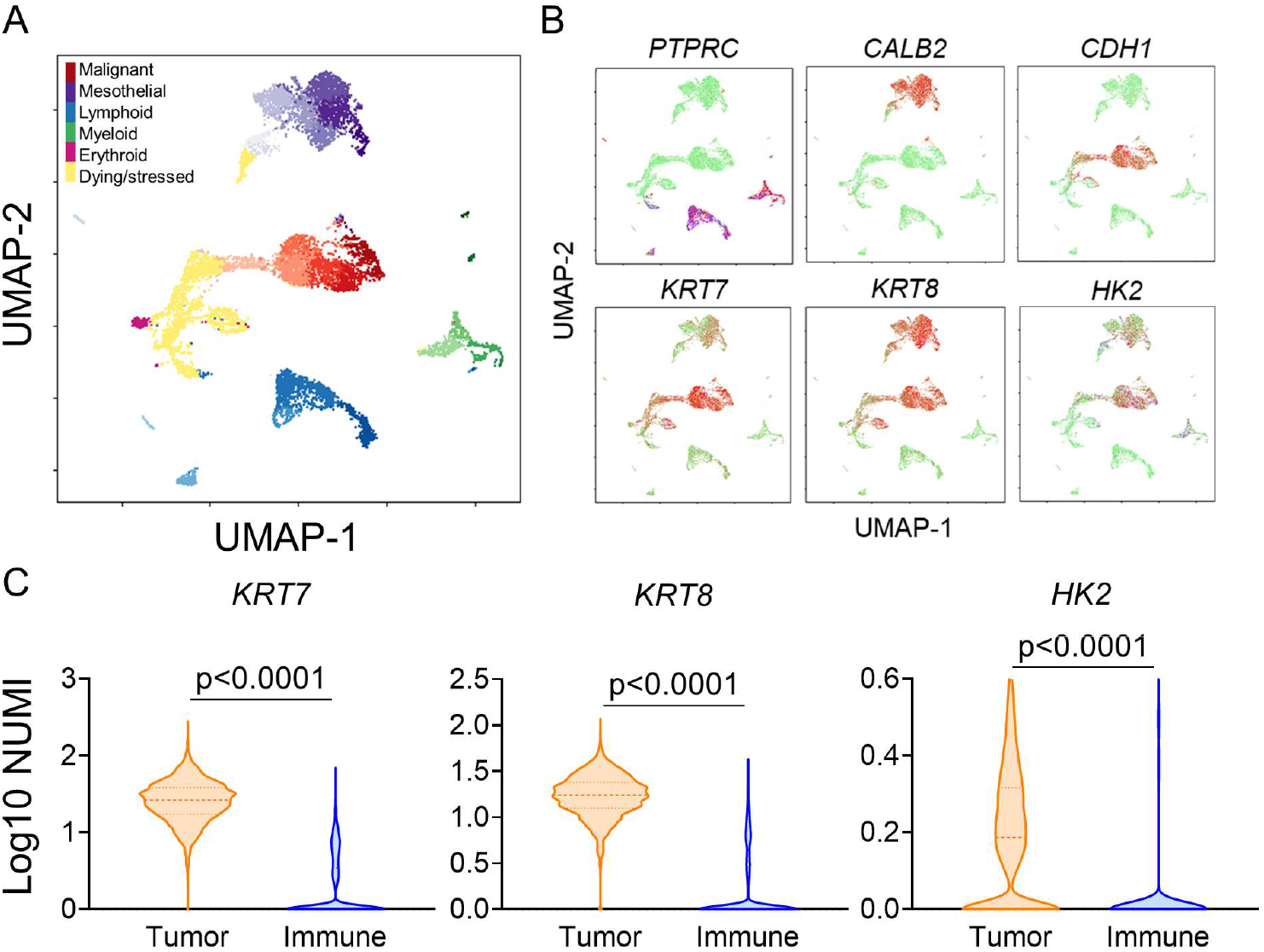
Single-cell transcriptome analysis of an MPE sample from an LUAD patient. (A) UMAP clustering of different cell types in the MPE sample. Different cell types are color-coded and separated at the different regions in the 2-dimensional space. The sub-clusters in each cell type cluster are labeled by the respective color with varying lightness. (B) Expression of selected marker genes across different cell types visualized in the 2-dimensional UMAP space. Green color denotes low expression and red color denotes high expression. (C) Comparison of KRT7, KRT8 and HK2 expression levels between CTCs and immune cells (N=2424 and 2009, respectively). The dash and dot lines of each violin plot denote the median and first and third quartiles, respectively.

HK2^high^/CK^neg^ CTCs were also detected in the CSF samples of LUAD patients. In 1.5 mL of CSF from P32, a total of 202 HK2^high^/CK^pos^/CD45^neg^ CTCs, 11 HK2^high^/CK^neg^/CD45^neg^ CTCs, and 77 HK2^low^/CK^pos^/CD45^neg^ CTCs were identified (Fig. 3D and table S3). Similar to MPE samples, both the CK^pos^ and CK^neg^ phenotypes exhibited consistent CNV profiles at the single-cell level (Fig. 4E). Across the 3 CSF samples measured, HK2^high^/CK^neg^ CTCs constituted a minority population in the CSF, ranging from 0.44% to 3.8% (Fig. 3C and table S3). Similar to the CTCs found in blood samples, morphological evaluation of CTCs in MPE and CSF samples showed that CK^pos^ CTCs are larger than CK^neg^ CTCs and this property may be attributed to the loss of cytokeratins within the CK^neg^ cells (Fig. 3F)

### Connection between CK expression and epithelial-mesenchymal transition

Different members of the CK family are common epithelial markers used in cytopathological analysis. Reduced CK expression in cancer cells is often associated with loss of epithelial signatures and EMT processes (*21*). To verify whether CK^neg^ CTCs acquired mesenchymal features and thus became more invasive, we performed single-cell RNA sequencing of an MPE sample collected from an LUAD patient, using 10X Genomics. A dimension reduction of the single-cell data using the Uniform Manifold Approximation and Projection (UMAP) algorithm clearly segregated the immune cells, mesothelial cells, and putative CTCs into different clusters, confirmed by the respective marker expression profiles (Fig. 4, A and B) (*22*). Specifically, the immune cell populations (blue and green clusters) showed clear CD45 expression. The cells in the upper purple cluster exhibited high calretinin expression and loss of E-cadherin – a marker combination that is specific to reactive mesothelial cells in cytological specimens (Fig. 4B) (*23, 24*). In contrast, most cells in the central red cluster display high E-Cadherin and CK7/8 expression without CD45 and mesothelial marker calretinin, suggesting a high likelihood of malignant cells (Fig. 4, A and B). Dying and stressed cells, as identified by the yellow cluster, exhibited extensive mitochondrial gene contamination and were removed from the subsequent analyses (Fig. 4A and fig. S5, see Materials and Methods).

To further verify that the cells in the red cluster are indeed tumor-derived cells, we employed an expectation-maximization algorithm to infer the CNV of all viable cells using single-cell transcriptome data as input (see Materials and Methods) (*25*). The putative CTCs showed distinct CNV patterns compared to normal cells. A k-mean clustering of the inferred single-cell CNV profiles clearly separated all the putative CTCs in the red cluster from the non-malignant mesothelial and immune cells in the other clusters, with no exception (fig. S6). Compared with the baseline CNV profile of normal cells, CTCs showed drastically different CNV profiles across many chromosomal arms, including clear amplification in chromosomes 5p and 14q as well as deletion in chromosomes 13q and 17p. These results confirmed the tumor origin of the putative CTCs identified by the UMAP algorithm. We further assessed the expression levels of HK2, CK7 and CK8 across single cells. A large number of CTCs showed significantly higher expression levels of HK2, CK7, and CK8 than immune cells (Fig. 4C). These results are consistent with the immunostaining results of the MPE samples, where the majority of the CTCs are CK^pos^.

CK7 and CK8 are highly expressed epithelial markers in NSCLC and are commonly used in cytopathological analysis of LUAD tumors (*26, 27*). Reduced CK expression in cancer cells is often associated with loss of epithelial signatures and EMT processes (*21*). To interrogate whether the CK^neg^ CTCs are those undergoing EMT and acquiring mesenchymal phenotype, we first defined a quantitative EMT score using the hallmark EMT gene set that contains 200 EMT-defining genes curated in the molecular signature database (MSigDB, Broad Institute) (*28*). The location of each single CTC in the EMT spectrum was quantified by the respective enrichment score obtained in the gene set enrichment analysis (GSEA) with respect to an average reference CTC (see Materials and Methods). Positive enrichment scores denote acquisition of mesenchymal signature and negative scores represent epithelial phenotype. A total of 426 cells and 58 cells were identified to have statistically significant epithelial and mesenchymal signatures (FDR<0.05), respectively, with many other cells sitting in the intermediate states of the EMT spectrum (Fig. 5A).

**Figure 5.**
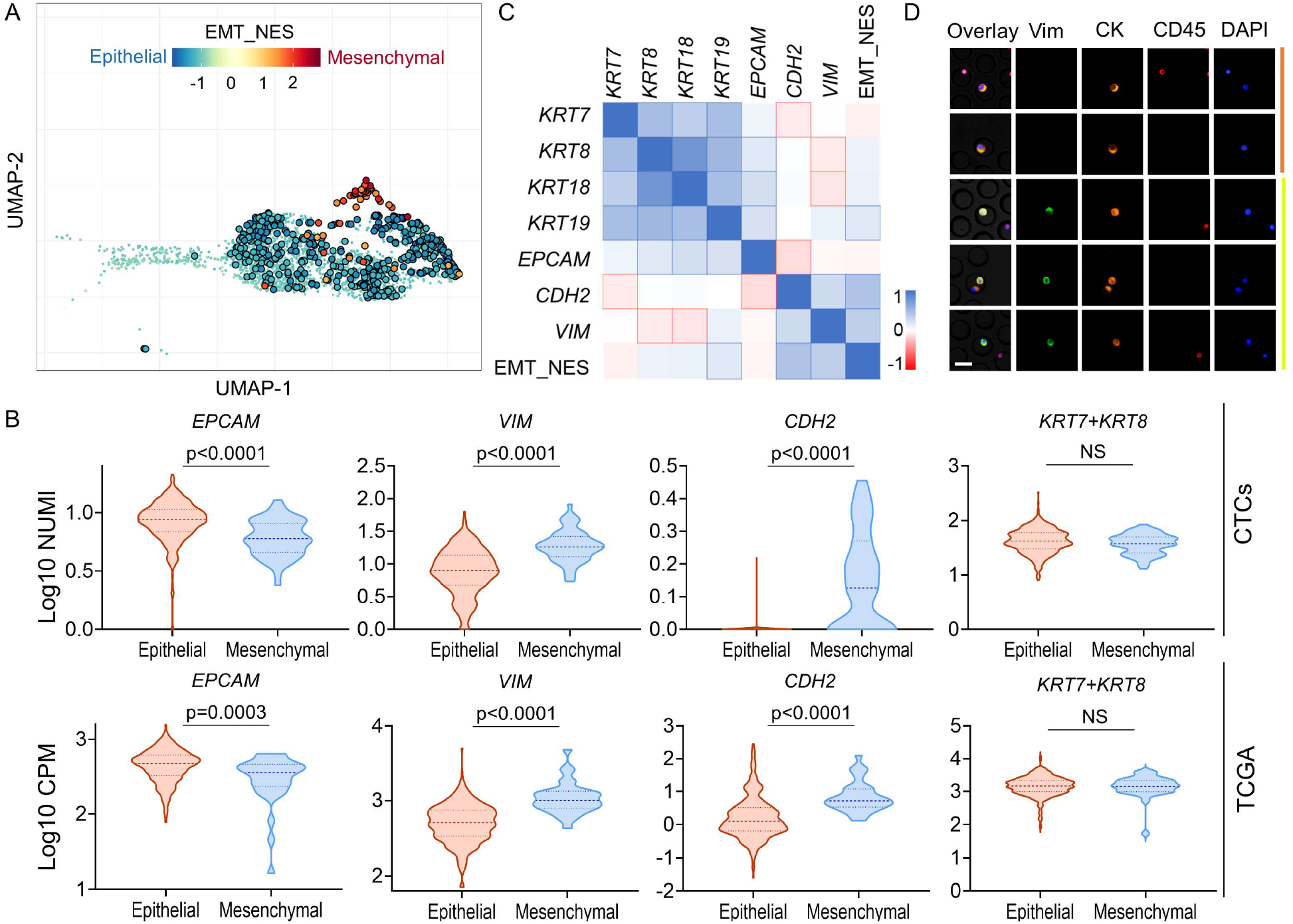
Molecular analysis of the CTCs with enriched EMT features. (A) Enrichment of EMT features of all the CTCs shown in the 2-dimensional UMAP space. The normalized enrichment scores (NES) are color-coded and cells with statistically significant enrichment scores (FDR<0.05) are highlighted by enlarged circles. (B) Comparison of expression levels of several EMT-related genes between epithelial and mesenchymal CTCs (top, N=426 and 58, respectively) and between epithelial and mesenchymal LUAD patient samples curated in TCGA (bottom, N=349 and 32, respectively) (NS: not significant). The dash and dot lines of each violin plot denote the median and first and third quartiles, respectively. (C) Spearman correlations between relevant cytokeratin family genes and EMT-related genes. Correlation coefficients are color-coded and statistically significant correlations (Bonferroni corrected p<0.05) are highlighted by outlines. (D) Representative fluorescence images showing the existence of both CK^pos^/Vimentin^pos^ and CK^pos^/Vimentin^neg^ CTCs in patient blood samples.

In line with the classifications described above, mesenchymal cells with high EMT scores show statistically lower *EPCAM* expression as well as higher *VIM* and *CDH2* expression than their epithelial counterparts (Fig. 5B). However, to our surprise, the difference of CK7/8 (*KRT7*+*KRT8*) expression levels between mesenchymal and epithelial cells is marginal (Fig. 5B). This lack of a clear difference in expression of these markers also holds true for other two types of highly expressed cytokeratin, *KRT18* and *KRT19*, in LUAD cells (fig. S7). The indistinguishable CK expression levels between epithelial and mesenchymal CTCs are consistent with the results that only weak or insignificant correlations were observed between various types of CK and EMT scores (Fig. 5C). In line with these results in gene expression levels, we found both CK^pos^/VIM^pos^ and CK^pos^/VIM^neg^ CTCs in patient blood samples using immunostaining, implicating the existence of CK^pos^ cells in both mesenchymal and epithelial CTCs (Fig. 5D).

These unexpected results between the CK expression and cellular EMT status in CTCs prompted us to interrogate whether differential expression of CK7/8 between mesenchymal and epithelial phenotypes exists in patient tumor tissues. We evaluated the gene expression data of LUAD patient samples curated in The Tumor Genome Atlas (TCGA) database, which includes 533 samples from primary tumor tissues and 2 from recurrent tumors. To classify their phenotypes in the EMT spectrum, we calculated the EMT scores using the analysis we employed for CTCs above (see Materials and Methods). Consistently, no statistically significant difference was identified in CK7/8 (*KRT7+KRT8*) expression levels between samples with epithelial signatures and those with mesenchymal signatures (FDR<0.05), while significant differences were observed in the expression levels of other EMT markers, including *EPCAM*, *VIM*, and *CDH2* (Fig. 5B). Taken together, these results suggest that the expression levels of CK7/8 – commonly used epithelial markers in cytopathological staining of LUAD – cannot effectively distinguish epithelial and mesenchymal CTC phenotypes in LUAD. This conclusion is also true for two other highly expressed cytokeratins (CK18/19) in lung cancer (fig. S7).

### Molecular signatures associated with CK^high^ and CK^low^ populations

The distribution of CK7/8 expression in epithelial and mesenchymal CTC populations is largely overlapping (Fig. 5B). Both populations have a few outlier CTCs with high (or low) CK expression levels and a large number of cells with intermediate CK levels. To interrogate the differential transcriptome profiles between CK-outlier CTCs in each phenotype, we first ranked all the CTCs based on their CK7/8 expression levels. We then denoted the cells within top 10% and bottom 10% of CK7/8 expression levels across all the CTCs as CK^high^ and CK^low^, respectively, and identified differentially expressed genes (DEGs) between them (see Materials and Methods).

A survey of the top-100 DEGs upregulated in CK^high^ cells revealed a number of cytokeratins and S100 family proteins shared by two phenotypic populations (Fig. 6A and fig. S8). CK^high^ cells displayed elevated γ-synuclein (*SNCG*) compared to CK^low^ counterparts in the mesenchymal population. Increased expression of γ-synuclein has been implicated in the pathogenesis of NSCLC by promoting cell survival and proliferation (*29*). Consistently, increased expression of proliferating cell nuclear antigen (*PCNA*) was also observed in CK^high^ cells in the mesenchymal population. In contrast, CK^high^ cells in the epithelial population were found to have increased expression in genes encoding integrin subunits (*ITGA2, ITGA3*, etc.) and MHC-I (*HLA-A, HLA-B, HLA-C*). For CK^low^ cells the top-100 upregulated DEGs included many genes encoding 40S or 60S ribosome protein components (*RPS2, RPL18*, etc.) as well as pulmonary surfactant-associated proteins (*SFTPA1, SFTPB,* etc.), which are either unique to a single phenotypic population or shared between two populations (Fig. 6B and fig. S8). Overexpression of genes encoding ribosome protein components in CTCs has recently been shown to contribute to tumor metastasis (*30*). Consistently, in CK^low^ cells of the mesenchymal population, we also found increased expression of *LOX* and *JUNB,* which have been implicated in enhanced invasiveness and tumor metastasis (*31–33*).

**Figure 6.**
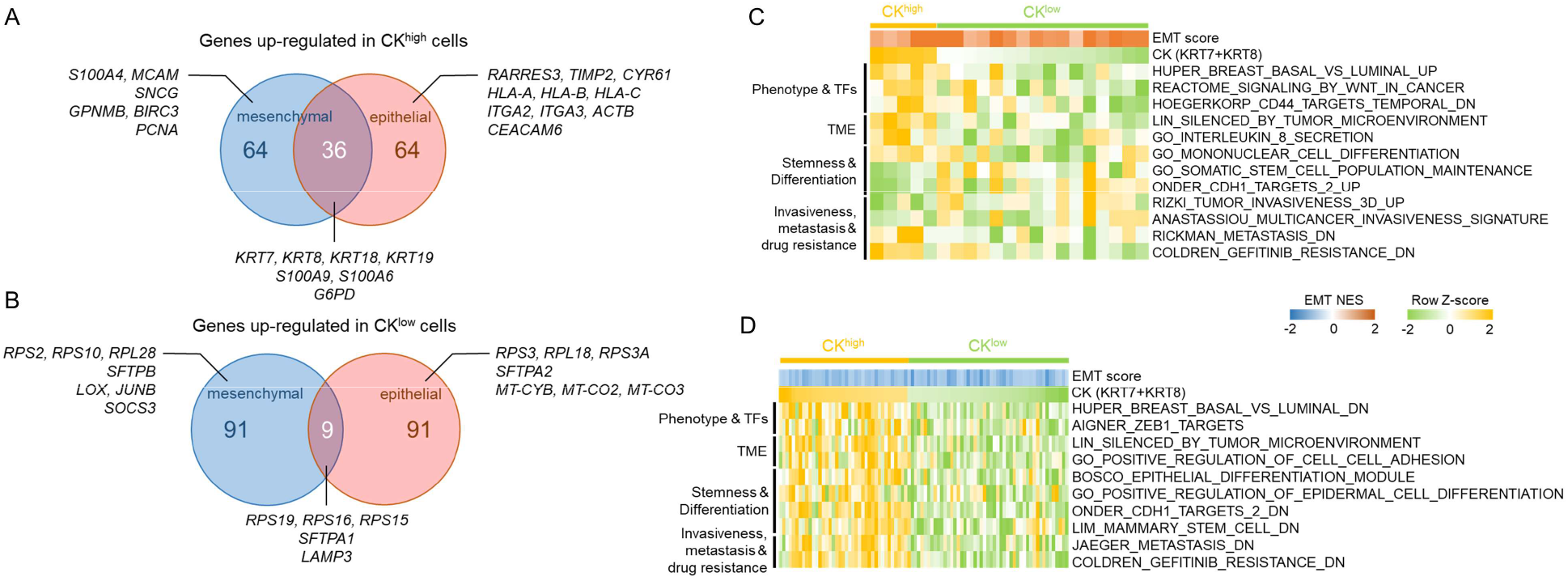
Molecular signatures associated with CK^high^ and CK^low^ CTCs. (A) Venn diagram showing the top-100 DEGs between CK^high^ and CK^low^ CTCs upregulated in CK^high^ cells shared between mesenchymal and epithelial populations as well as specific to each population. (B) Venn diagram showing the top-100 DEGs between CK^high^ and CK^low^ CTCs upregulated in CK^low^ cells shared between mesenchymal and epithelial populations as well as specific to each population. (C) Enrichment scores of selected gene sets in CK^high^ and CK^low^ CTCs in the mesenchymal population analyzed by GSVA. (D) Enrichment scores of selected gene sets in CK^high^ and CK^low^ CTCs in the epithelial population analyzed by GSVA.

To resolve the molecular programs associated with CK^high^ and CK^low^ cells in both phenotypic populations, we performed gene set variation analysis (GSVA) across all the CK-outlier CTCs in each phenotype (*34*). We interrogated the enrichment profiles of selected gene sets that displayed statistically significant differences between CK^high^ and CK^low^ cells (Fig. 6, C and D). Among these gene sets, CK^low^ cells showed enrichment in stem cell features and target genes in response to CDH1 knockdown in both mesenchymal and epithelial populations (Onder_CDH1_targets_2) (*35*). These cells also displayed signatures associated with metastasis and EGFR tyrosine kinase inhibitor resistance, as evidenced by the enrichment profiles in several relevant gene sets. For example, most CK^low^ cells in the epithelial population have negative enrichment scores and consequently reduced expression levels in the Jaeger_Metastasis_DN gene set that includes genes down-regulated in metastatic tumors (Fig. 6D) (*36*). Similarly, a majority of CK^low^ cells in the mesenchymal population also displayed negative enrichment scores in a set of genes down-regulated in metastatic tumors (Fig. 6C). Taken together, both the DEG and GSVA analyses suggest that CK^low^ cells exhibit metastasis-related molecular signatures compared to their CK^high^ counterparts, regardless of their EMT status.

## Discussion

Cancer cells often elevate glucose metabolism in order to fuel their uncontrolled growth. Exploiting this altered metabolism allowed us to develop metabolic activity-based methods for rapid identification and retrieval of CTCs from liquid biopsy samples for both epithelial and nonepithelial malignancies (*37*). In the studies presented here, we evaluated a key enzyme in glucose metabolism – HK2 – as a surrogate for metabolic activity-based CTC detection. HK2 catalyzes glucose phosphorylation, the first enzymatic step in glycolysis, and is expressed at easily detectable levels in many cancer cells. The use of HK2 as a marker allows us to analyze fixed blood cells preserved in TransFix/EDTA vacuum blood collection tubes within 3 days of blood collection, without significant signal loss. This approach relieves the time constraint and pressure of sample delivery, processing and metabolic analysis, and thus significantly improves the adaptability of our assay. Single-cell sequencing confirmed the malignancy of HK2-identified putative CTCs. All the randomly selected HK2^high^ cells from different types of liquid biopsy samples were confirmed to be malignant by single-cell genome-wide CNV analysis (Figs. 2D; 3, B and E, and figs. S3 and S4), attesting the high specificity of HK2 as a glycolytic activity-associated marker in identifying CTCs present in a high background of leukocytes and other confounding cell types.

The use of HK2 as a tumor cell marker permits us to reveal the CK^neg^ subpopulation of CTCs, a population of CTCs which is not accessible to current epithelial marker-based CTC detection methods. Although new markers (e.g. oncofetal chondrotin sulfate) or a combination of multiple markers have been developed to capture multiple phenotypes of CTCs (*38, 39*), these methods are still dependent on CK expression and thereby inherently biased toward identifying CK^pos^ CTCs. In liquid biopsy samples, the existence of CK^neg^ malignant cells are normally not easily identified, because many normal cell types with low or undetectable CK expression levels confound such examination. Sequencing all the CK^neg^ cells in a liquid biopsy sample at the single-cell level to identify CK^neg^ CTCs is neither practical nor cost-effective. Additionally, we quantitatively evaluated the EMT status of each CTC based upon its overall transcriptional enrichment in 200 EMT-defining genes rather than relying on single or a few EMT markers (Fig. 5A). Our results indicate that CK expression levels are largely independent of cellular EMT status in LUAD. Both epithelial and mesenchymal CTC populations contain cells with high (or low) CK expression. Consistently, both CK^pos^/VIM^pos^ and CK^pos^/VIM^neg^ CTCs can be found in patient blood samples (Fig. 5D). The cells with low CK expression in the epithelial population normally do not express mesenchymal markers. Therefore, inclusion of mesenchymal markers, such as vimentin and N-cadherin, in the detection panel would not necessarily identify CK^neg^ CTCs. HK2 and CD45 co-staining can distinguish CTCs from a large number of immune cells and other confounding cell types in liquid biopsy samples and allows identification of both CK^pos^ and CK^neg^ CTCs.

HK2^high^/CK^neg^ CTCs were found to be a prevalent CTC subtype in 50% of blood samples from 16 LUAD patients with detectable CTCs (Fig. 2E). Six of these samples contained only CK^neg^ CTCs and standard EpCAM/CK-based CTC detection methods are unable to detect them. This observation partially explains why CTCs are less frequently observed in peripheral blood of NSCLC patients compared to other epithelial cancers when detected by CellSearch-like systems. Some patients have predominantly HK2^low^/CK^pos^ CTC population in their blood, as exemplified by P2, which may be attributed to a quiescent or stressed cell state with reduced glycolysis and HK2 expression (*40*). Our approach does not detect those CK^neg^ CTCs with low HK2 expression, because of contaminating circulating cells of normal tissue or blood origin. Nevertheless, we detected 5 or more CTCs in 5 mL blood samples from 37.5% of NSCLC patients, including two stage III patients with localized disease. This level of sensitivity is clearly greater than that of previous reports, where 5 or more CTCs were detected in 7.5 mL blood from only 15% of stage IV NSCLC patients and no patients with localized disease having more than 4 CTCs (*6*).

To our knowledge, this study is the first characterization of CTCs from multiple liquid biopsy forms of the same tumor type. HK2^high^/CK^neg^ cells were a prevalent CTC subtype in 50% of blood samples with positive CTC counts but were rarely detected in MPE and CSF of LUAD patients. The differences in CK subpopulations across different types of liquid biopsy may be the consequences of different microenvironments of these liquid biopsy samples and/or distinct mechanism of dissemination between CK^pos^ and CK^neg^ CTCs. While the expression of CK was not correlated with cellular EMT status, cells with low CK expression were found to be enriched in more metastasis-related transcriptional signatures compared to those with high CK expression (Fig. 6). Thus enrichment of CK^neg^ CTCs in peripheral blood samples might result from the CTCs shedding from metastatic lesions. An alternative hypothesis would be that CTCs in the blood might tend to transition to a CK^neg^ subtype, acquire a more metastatic signature than those in MPE or CSF and, consequently, seed distant organs. In all types of liquid biopsy samples, CK^neg^ CTCs have smaller sizes than CK^pos^ CTCs, possibly due to the reduced expression of intermediate filaments (Figs. 2A and 3F). However, their CNV profiles resemble those of CK^pos^ CTCs, suggesting that CK expression levels are likely to be epigenetically regulated rather than reflecting the existence of genetically distinct clones (Figs. 2D; 3, B and E) (*41*). The phenotypic transitions between CK^pos^ and CK^neg^ CTCs, as well as the clinical implications of CK^neg^ CTCs as a prognostic marker or a drug target in LUAD patients, warrant further investigation.

## Materials and Methods

### Cell lines and reagents

Hep3B, HepG2, H460, HCC827, H1975, H1299, HT29, HCT116, SW620, and Caco2 cells were from the American Type Culture Collection (ATCC). Huh7, JHH7, JHH5, and MDAMB468 cells were provided by Drs. Dennis Slamon and Richard Finn (UCLA). SUM159, MDAMB231, and HCC1937 cells were provided by Dr. Heather Christofk (UCLA). A549, SKOV3, OVCAR5, and HEY cells were provided by Dr. Caius Radu (UCLA). Hs766T, MiaPaca2, and Panc1were provided by Dr. Timothy Donahue (UCLA). LNCaP, DU145, and PC3 cells were provided by Dr. Hong Wu (UCLA). 786-0 cells were provided by Dr. Allan Pantuck (UCLA). All cells used for the experiments were between passages 3 and 20. Cell lines were routinely authenticated based on morphology and growth characteristics. All cells were maintained in RPMI-1640 Medium (Life Technologies) containing 1× Penicillin-Streptomycin-Glutamine (Gibco) and 10% FBS (Gibco) in humidified atmosphere of 5% CO_2_ and 95% air at 37°C. Cells were routinely checked for Mycoplasma contamination by using MycoAlert (Lonza). (APC)-conjugated CD45 (clone HI30), Alexa Fluor 488-conjugated goat-anti-rabbit secondary antibody (#A11008) and MitoTracker™ Green FM (#M7514) were purchased from ThermoFisher Scientific. PE-conjugated anti-cytokeratin 7/8 antibody was purchased from BD Biosciences (#347204). Anti-HK2 primary antibody was purchased from Abcam (#ab209847). Anti-Vimentin antibody was purchased from CST (#5741S). DAPI was purchased from Beyotime Biotechnology. Hank’s balanced salt solution (HBSS, no calcium, no magnesium, no phenol red) was purchased from Gibco. All primers were synthesized by Genewiz and listed in Table S4.

### Fabrication of microwell chip

The microwell chip was fabricated in poly(dimethylsiloxane) (PDMS) using standard microfabrication soft-lithographic techniques. A replicate for molding the PDMS was obtained by patterning a silicon wafer using photoresist SU-8 2050. The PDMS pre-polymer (Sylgard 184, Dow Corning) was mixed in a ratio of 10:1, and subsequently casted on this lithographically patterned replicate. After curing at 80 °C for 2 h, the PDMS component was separated from the replicate. The diameter and depth of the microwells are 30 μm and 20 μm, respectively. A chip containing 28,000 microwells was used to measure peripheral blood and cerebrospinal fluid samples, and a chip containing 112,000 microwells was used to measure pleural effusion samples.

### Patient information and sample collection

Peripheral blood and pleural effusion samples of lung adenocarcinoma were obtained from patients in Shanghai Chest Hospital with written informed consents. Cerebrospinal fluid samples were obtained from lung adenocarcinoma patients in Huashan Hospital with written informed consents. The clinical sample collection and experiments were carried out in accordance with guidelines and protocols that approved by the Ethics and Scientific Committees of Shanghai Chest Hospital and Huashan Hospital. The tumors were staged according to the 7th edition of the international tumor-node-metastasis (TNM) system.

### Identification of CTCs in peripheral blood samples

Fresh blood samples (5 mL) were drawn and preserved in TransFix/EDTA Vacuum Blood Collection Tubes and delivered to the lab within 4 hours. Blood samples were initially centrifuged at 500 g for 5min. The supernatant was discarded and the cell pellet was re-suspended in an equivalent volume of HBSS and mixed with 25μL CTC enrichment antibody cocktail (RosetteSep™ CTC Enrichment Cocktail Containing Anti-CD36, STEMCELL Technologies) at room temperature for 20 min. 15 mL HBSS with 2% FBS (Gibco) was then added and the samples were mixed well. The mixture was carefully added along the wall of Sepmate tube (SepMate™-50, STEMCELL Technologies) after adding 15 mL density gradient liquid (Lymphoprep™, STEMCELL Technologies) into the tube through the middle hole. After centrifuging at 1200 g for 20 min, the topmost supernatant (~10 mL) was discarded, and the remaining liquid (~10 mL) above the barrier of the Sepmate tube was rapidly poured out into a new centrifuge tube. After centrifuging at 600 g for 8 min, the supernatant was removed and 1 mL of red blood cell lysing buffer (BD Biosciences) was then added for 5 min to lyse red blood cells. After centrifuging at 250 g for 5 min, the nucleated cell pellet was re-suspended in HBSS. The cell suspension was then applied onto the 3% BSA (Sigma)-treated microwell chip. A 10 min waiting period was allowed for the cells to sit down in the 28,000 microwells. The chip was sealed with a porous Isopore™ polycarbonate membrane (Merck Millipore, #TSTP04700) to avoid cell loss during subsequent on-chip staining. After cell fixation (2% PFA, 10 min) and permeabilization (0.5% Triton X-100, 15 min), blocking solution consisting of 3% BSA and 10% Normal Goat Serum was applied to the chip for 1 h, followed by incubation with APC-conjugated anti-CD45 antibody (mouse), PE-conjugated anti-CK 7/8 antibody (mouse) and anti-HK2 antibody (rabbit) in PBS overnight at 4°C. After extensive washing with PBS, cells on the chip were treated with Alexa Fluor 488-conjugated goat-anti-rabbit secondary antibody in PBS for 1 h and DAPI for 10 min followed by washing with PBS. ImageXpress Micro XLS Widefield High Content Screening System (Molecular Devices) scanned the chip and imaged all cells in bright field and four fluorescent colors (CD45: CY5; HK2: FITC; CK: TRITC; Nucleus: DAPI). A computational algorithm analyzed the images and identified HK2^high^ and CK^pos^ cells based on the cut-offs generated from HK2 and CK fluorescence intensity of CD45^pos^ leukocytes in the samples.

### Identification of CTCs in malignant pleural effusion (MPE) and cerebrospinal fluid (CSF) samples

For CTC identification in MPE, 5 mL of an MPE sample was first filtered by a membrane with a pore size around 100 μm, followed by centrifuging at 500 g for 5 min to separate cell pellets. 1 mL of red blood cell lysis buffer (BD) was then added to lyse red blood cells for 5 min. After centrifuging at 500 g for 5 min, the nucleated cell pellet was re-suspended in and washed with HBSS. After cell counting, up to 500,000 cells were applied onto an 112,000-well microwell chip for cell fixation, permeabilization and immunostaining as described above. For CTC identification in CSF, CSF samples were centrifuged at 500 g for 5 min to separate cell pellets and re-suspended in HBSS, followed by processing with the microwell chip. No enrichment step was included, due to the limited number of cells present in the CSF samples.

### Single-cell manipulation and *EGFR* mutation detection

Individual candidate CTCs in the microwells were retrieved using a XenoWorks Micromanipulator and trimethylchlorosilane (TMCS)-treated micropipettes, and then transferred into PCR tubes (Axygen). The genome amplification of the retrieved single cells was conducted with the MALBAC® Single Cell Whole Genome Amplification (WGA) Kit (Yikon Genomics). For detection of *EGFR* L858R mutation, PCR reaction was performed using the primers listed in table S4. The PCR reaction buffer consisted of 12.5μL 2×Ex Taq DNA polymerase mix (VazymeBiotech), 10μM forward primer, 10 μM reverse primer and 0.2μL whole genome amplified DNA. The PCR reaction was conducted as follows: 95°C for 3 min, followed by 30 cycles (95°C for 30 s, 60°C for 30 s and 72°C for 30s), followed by a final extension at 72°C for 5min. The PCR products were analyzed by Sanger sequencing (Genewiz, Suzhou, China).

### Single-cell whole genome sequencing

We used 22 pairs of primers targeting 22 loci on different chromosomes to evaluate the WGA coverage of single cells (see table S4 for primer sequence design) (*37*). WGA products with 18 out of 22 loci amplified were used for subsequent whole genome sequencing (WGS) library construction. For library construction, WGA products were firstly digested with dsDNA fragmentase (New England Biolabs) and 300-500bp fragments were retrieved with Agencourt AMPure XP Beads (Beckman Coulter). Around 100 ng of DNA fragments were used as input for sequencing library construction. DNA fragment repair and library adaptor ligation were performed using NEBNext® Ultra™ DNA Library Prep Kit for Illumina (New England Biolabs), in accordance with the manufacturer’s protocol. WGS libraries were amplified using NEBNext® Ultra™ II Q5® Master Mix and index primers (New England Biolabs). Library purification was performed with TIANgel Midi Purification Kits (TIANGEN Biotech). The concentrations of purified fragmented DNA or libraries were quantified with Qubit dsDNA HS Assay Kits (Invitrogen). Libraries were analyzed by Illumina HiSeq X Ten platform with 150 PE at 0.1X coverage (Genewiz, Suzhou, China).

### Copy number determination and segmentation from whole genome sequencing data

Sequencing reads were aligned to the major chromosomes of human (hg19) using BWA (version 0.7.10-r806) with default options (*42*). SAMtools (version 1.31) was used to mark and remove PCR duplicates (*43*). To reduce WGS biases, the sequence depths of tumor cells were normalized by GC contents and mappability with 500-kb windows. The diploid regions were determined using HMMcopy (*44*). Similar copy numbers in adjacent chromosome regions were merged in DNAcopy package to get CNV regions (*45*).

### Processing 10X Genomics single-cell RNA sequencing data

Cells were collected from an MPE sample of a LUAD patient. Red blood cell lysis and leukocyte depletion (EasySep Human CD45 Depletion Kit II #17898, STEMCELL Technologies) of MPE were performed before processing with 10X Genomics. RNA-seq profiles of 10,026 single cells were obtained using the Cell Ranger pipelines (version 3.1.0) and the provided reference (refdata-cellranger-GRCh38-3.0.0). Clusters of cells were identified using the k-means clustering algorithm as implemented in the Cell Ranger pipeline. Mitochondrial count percentage was calculated for each cell (fig. S5). A cluster was defined as stressed or dying if the average mitochondrial count percentage was more than 30%. One special cluster was close to dying clusters in the UMAP and the expressed genes number of that cluster was small, indicating that this cluster contained cells of poor quality (fig. S5). After removing these stressful/dying and poor-quality cells, five main cell types, including malignant, mesothelial, myeloid, lymphoid, and erythroid cells, were identified based on the clustering and the UMAP dimension reduction (Fig. 4A).

### CNV inference using the single-cell transcriptome data

For each of those 10,026 single cells from 10X Genomics RNA sequencing, we inferred an estimated CNV profile using R package CONICS over 38 chromosome arms (fig. S6), where the CNV profile was simulated by the posterior probabilities based on a two component Gaussian mixture model (*25*). We first unbiasedly used all 44 autosome arms to infer CNV profiles; however only 38 of these analyses were able to fit 2-component mixture models. The clustering results showed that all 10,026 single cells can be divided clearly into two clusters, one of which represented malignant cells whose inferred CNV profiles are drastically different from those of benign cell types in the other cluster (fig. S6).

### Calculation of EMT scores using single-cell transcriptome data

The Molecular Signatures Database (MSigDB v7.0, Broad Institute) was used to calculate EMT scores through gene set enrichment analysis (*28*). R package fgsea was applied to the gene set HALLMARK_EPITHELIAL_MESENCHYMAL_TRANSITION to calculate a normalized enrichment score for each CTC as its EMT score, with 1000 permutations and the rank list of genes generated as follows: Only those genes observed in at least one CTC were considered. A reference cell was created with expression of each gene as the average expression of this gene across all the tumor cells. To rank genes for each cell with respect to the reference cell, log2_Ratio_of_Classes was used. Only cells with FDR<0.05 were considered significantly enriched in either epithelial or mesenchymal phenotypes.

### Analysis of differentially expressed genes

For the 10,026 single cells from 10X Genomics RNA sequencing, we used the normalized UMI counts for their expression levels. The normalization scaled the raw UMI counts for each barcode by multiplying ratio of the median UMI sum of all barcodes to the sum of UMI counts of this barcode, assuring that all normalized UMI sums of barcodes have the same as that of the median barcode. After normalizing and log10-transformation of UMI counts, to be consistent with our pan-CK (anti-CK7/8) staining used for CTC identification, the CK (cytokeratin) level in each of tumor cells was defined as the sum of *KRT7* and *KRT8* expression levels in the cell. CK^low^ cells were defined as cells with CK expression level lower than the 10% probabilistic quantile, and CK^high^ cells were defined as cells with CK level higher than the 90% probabilistic quantile. Any CTC was considered to be mesenchymal if its EMT score (see above Calculation of EMT scores using single-cell transcriptome data) was positive and FDR < 0.05, and epithelial if its EMT score was negative and FDR < 0.05. In either epithelial or mesenchymal population, R package LIMMA was used to compare expressions of CK^high^ cells versus CK^low^ cells and to calculate the fold change and p value for each gene. Only those genes with p values < 0.05 were considered as differentially expressed genes between CK^high^ cells and CK^low^ cells. Top 100 genes up-regulated in CK^high^ cells and down-regulated in CK^high^ cells (i.e. up-regulated in CK^low^ cells) in terms of fold changes were respectively selected and compared in Figure 6.

### Gene set variation analysis

To compare the molecular signatures differentially enriched in CK^high^ cells or CK^low^ cells in either epithelial or mesenchymal population, R package GSVA was applied to the CK^high^ and CK^low^ cells to derive a GSVA enrichment score for each cell under a certain gene set (*34*). Gene sets obtained from the Molecular Signatures Database (MSigDB v7.0, Broad Institute) include the H (hallmark gene sets), C2 (curated gene sets) and C5 (GO gene sets). Normalized and log10-transformed UMI counts were input as the expression matrix. Since GSVA would discard genes with constant expression values across all cells, only those genes with their coefficients of variation (CVs) larger than 1 were selected. We obtained a very similar matrix of GSVA enrichment scores using all the genes as well. All other parameters were set to their default values, except for the sizes of gene sets. We set the minimum size (min.sz) as 10 and the maximum size (max.sz) as 900. After deriving the gene-set-by-cell matrix of GSVA enrichment scores, we used R package LIMMA to compare GSVA enrichment scores of CK^high^ cells with those of CK^low^ cells and to determine if a gene set had a significantly differential enrichment between CK^high^ and CK^low^ cells for a given gene set, via score difference and p-value.

### Analysis of LUAD RNA-seq profiles from TCGA

HTSeq-counts profiles of 533 primary LUAD tumor samples and 2 recurrent tumor samples were downloaded from the Genomic Data Commons (https://portal.gdc.cancer.gov/). Raw read counts were first normalized to counts per million (CPM) using R package edgeR. A reference profile was then created by averaging over all 535 samples. For each of these 535 samples, the EMT score was calculated via gene set enrichment analysis in the same way described above using R package fgsea and the hallmark EMT gene set from MSigDB. The reference profile was created with expression of each gene as the average expression of this gene across all LUAD samples. The sample was considered to be mesenchymal if its enrichment score was positive and FDR < 0.05, and epithelial if its enrichment score was negative and FDR < 0.05.

### Statistical analysis

Statistical analyses were performed using GraphPad PRISM 8 (GraphPad Software, Inc) and XLSTAT (Addinsoft, for correlation test) unless noted elsewhere. Statistical significance between two groups was compared using the two-tailed Mann-Whitney test with p < 0.05 as the significance threshold. Alpha level was corrected by Bonferroni correction when multiple groups were compared.

## Data availability

The single-cell sequencing data reported in this paper have been deposited in the ArrayExpress database (accession no. E-MTAB-8767).

## Author contribution

H.R.H., S.L., Q.S. and W.W. designed and supervised the study. Q.Z., Y.H., Z.L. and M.X. contributed clinical samples. L.Y., J.C., S.X., Z.W. and Y.T. performed the experiments. X.Y. and Y.D. performed the computational and bioinformatic analysis. X.Y., Q.Z., Y.H., D.Z., Q.S. and W.W. analyzed the data. Q.S. and W.W. wrote the paper. S.X., H.R.H., Q.S. and W.W. revised the manuscript.

