## supplemental information for "Resolving an underrepresented circulating tumor cell population in lung cancer enabled by Hexokinase 2 analysis"

[1]Shanghai Bone Tumor Institute and Department of Orthopedics, Shanghai General Hospital, Shanghai Jiao Tong University School of Medicine, Shanghai, China; [2]Institute for Systems Biology, Seattle, WA, USA; [3]Key Laboratory of Systems Biomedicine (Ministry of Education), Shanghai Center for Systems Biomedicine, Shanghai Jiao Tong University, Shanghai, China; [4] Department of Oncology, Huashan Hospital, Shanghai Medical College, Fudan University, Shanghai, China; [5]Department of Molecular and Medical Pharmacology, David Geffen School of Medicine, University of California, Los Angeles, CA, USA; [6]Shanghai Lung Cancer Center, Shanghai Chest Hospital, Shanghai Jiao Tong University, Shanghai, China; [7]MinhangBranch, Zhongshan Hospital, Fudan University, Shanghai, China; [8]Shanghai Key Laboratory of Medical Epigenetics, Institutes of Biomedical Sciences, Fudan University, Shanghai, China;; [9]InstituteofFudan-Minhang Academic Health System, Minhang Hospital, Fudan University, Shanghai, China.

I. Supplementary Figures S1-S8

II. Supplementary Tables S1-S4

III. Supplementary Datasets S1-S2

**Supplementary Figures**

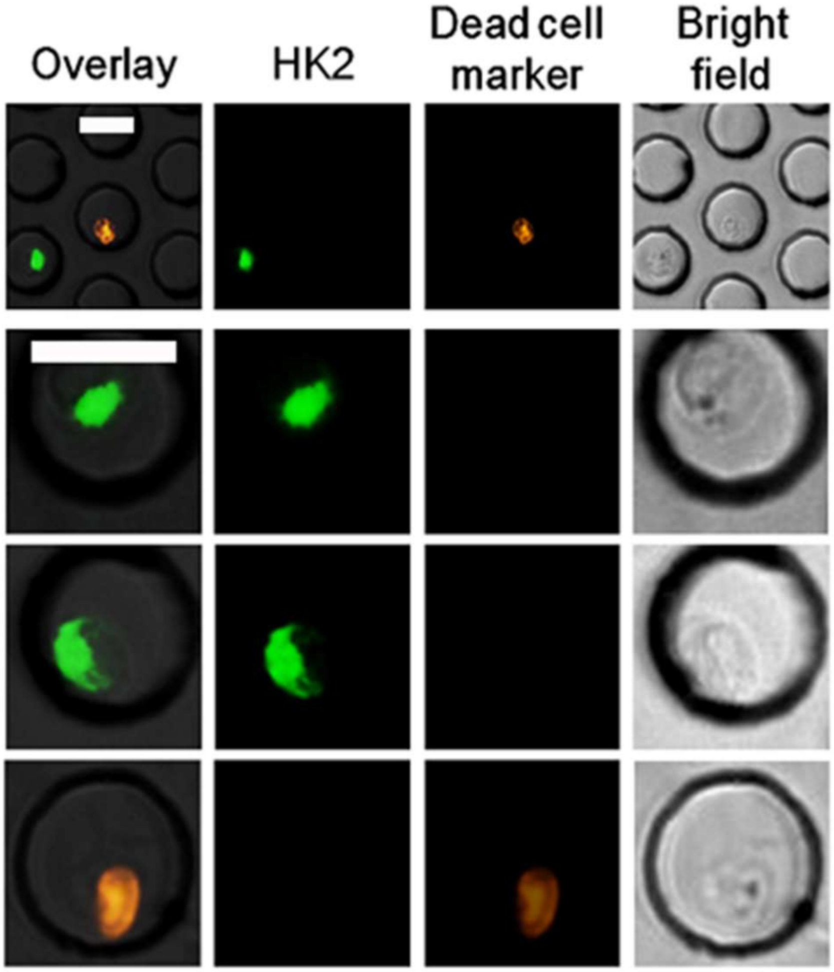


**Figure S1.** Dead H1975 lung cancer cells are absent of HK2 fluorescence signals. The dead cell marker is from the Live/Dead Viability/Cytotoxicity kit (Thermo Fisher, Catalog L3224). Scale bar: 30 μm.

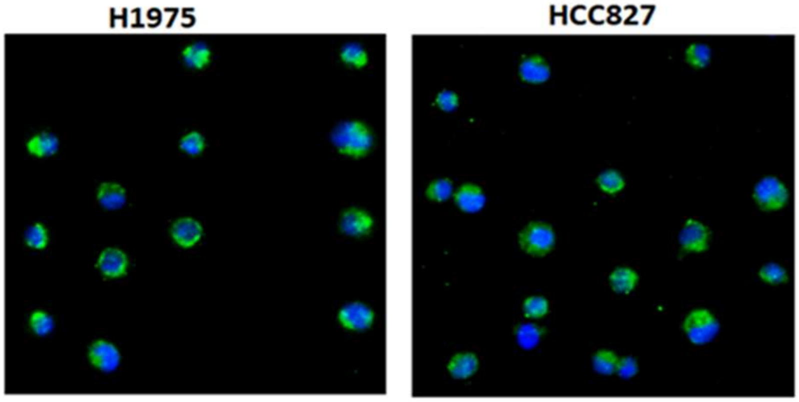


**Figure S2.** Representative fluorescence images of H1975 and HCC827 cells stained with anti-HK2 and DAPI.

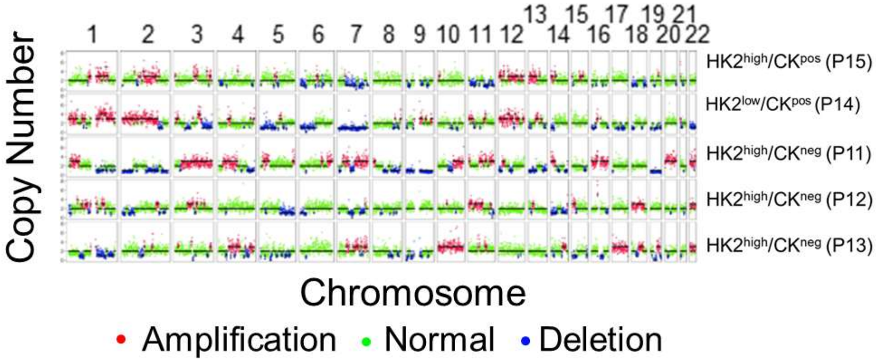


**Figure S3.** Single-cell CNV profiles across the chromosomes of randomly selected CTCs from five patients.

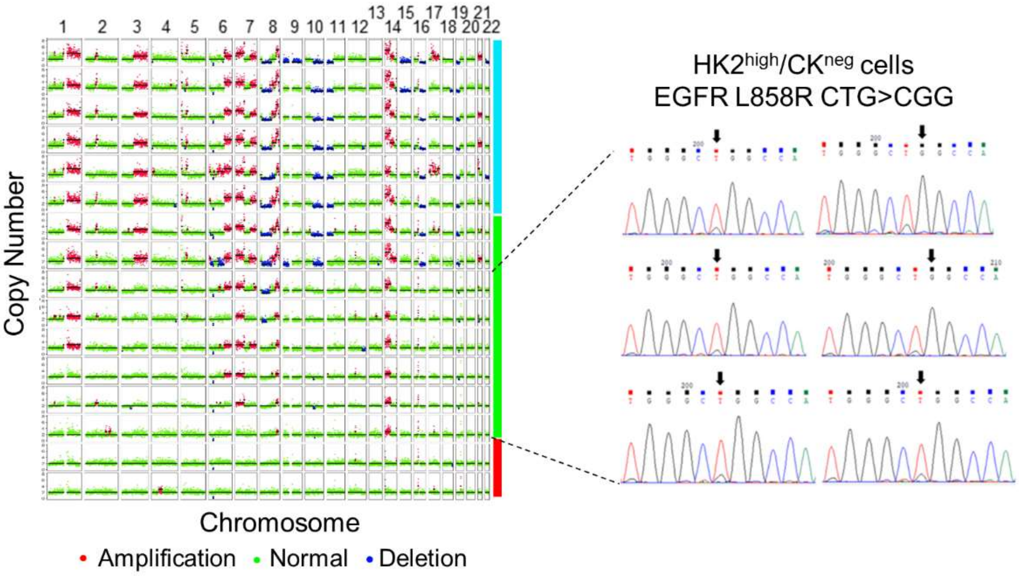


**Figure S4**. Single-cell CNV profiles of randomly selected CTCs and leukocytes from the MPE sample of P25. $HK2^{high}/CK^{pos}/CD45^{neg}$ and $HK2^{high}/CK^{neg}/CD45^{neg}$ CTCs as well as leukocytes are color-coded by cyan, green and red bars to the right, respectively. All the cells were detected to harbor *EGFR* L858R oncogenic driver mutation as their primary tumor lesion. The representative Sanger sequencing results of selected $HK2^{high}/CK^{neg}/CD45^{neg}$ CTCs are shown to the right.

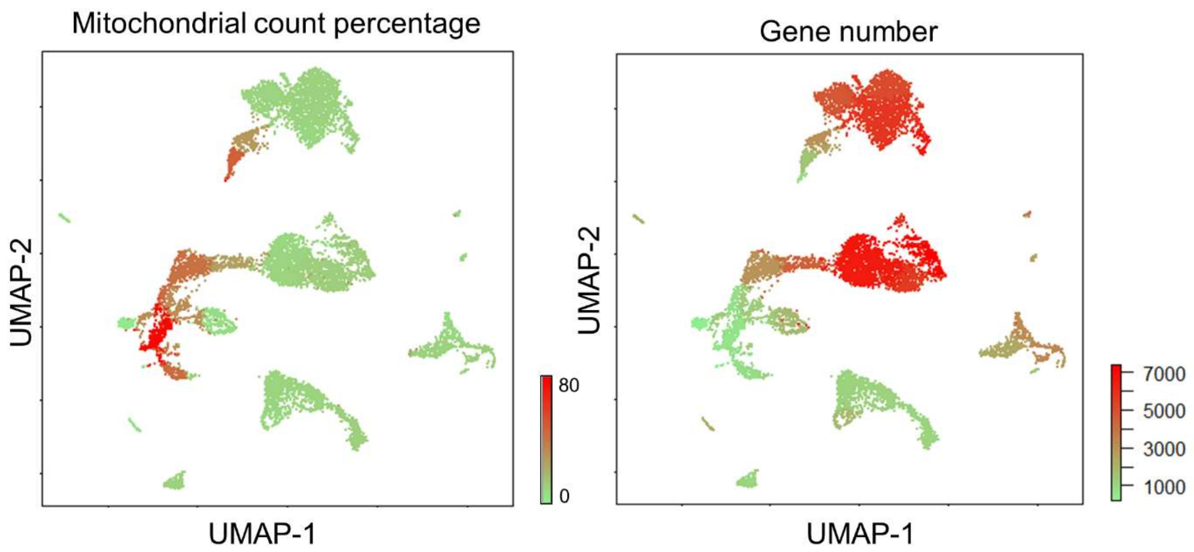


**Figure S5**. UMAP visualization of average mitochondrial count percentage (left) and gene number (right) of the cell clusters identified in Fig. 4A. Both of them are quality-control metrics commonly used to identify stressed and dying cells.

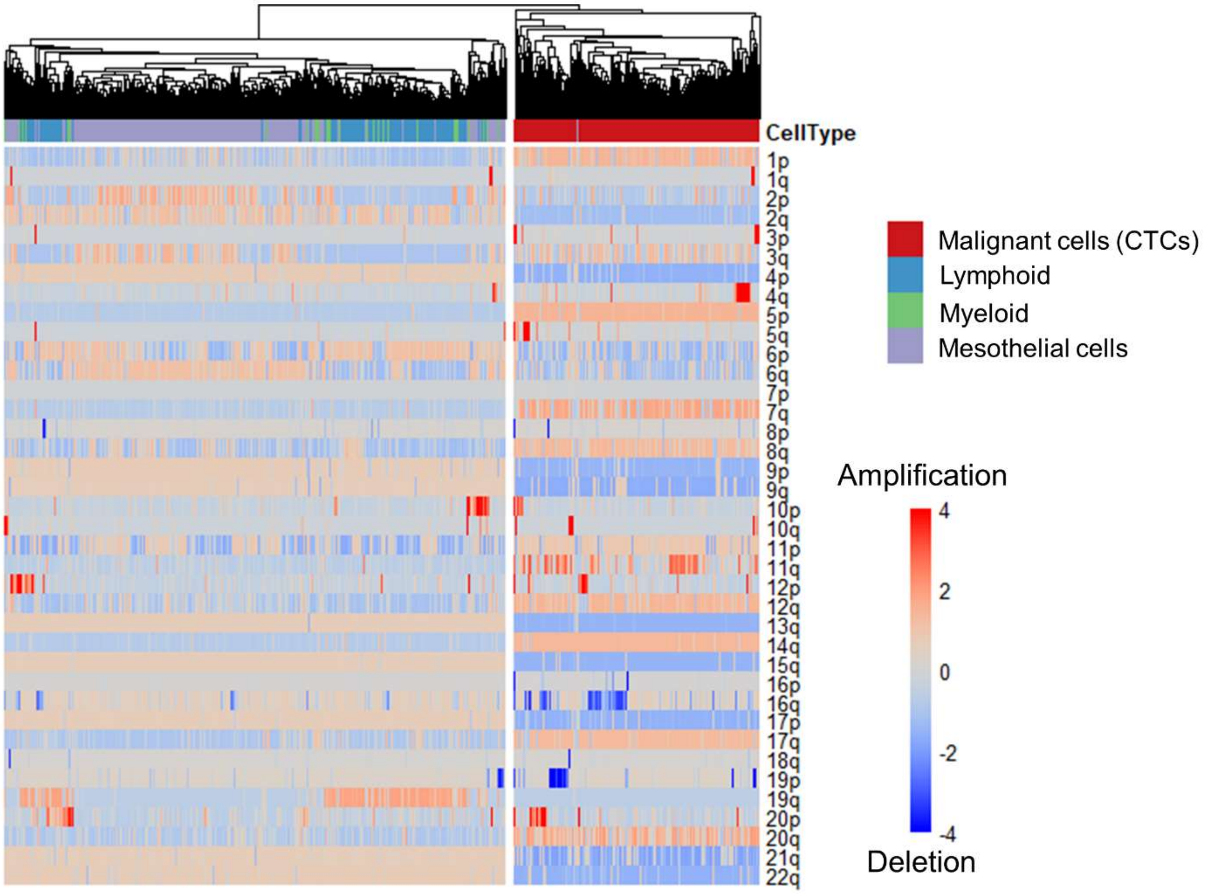


**Figure S6.** K-mean clustering of inferred single-cell CNV profiles across the chromosome. Chromosome arms that failed to fit CONICS two-component Gaussian mixture model were excluded. Cell types were labeled based on the UMAP clustering results in Fig. 4A. All the malignant cells (CTCs) are clustered to the right with significantly altered CNV profiles across the chromosome compared to the non-malignant cells in the left cluster. Stressed/dying and erythroid cells are excluded from the analysis.

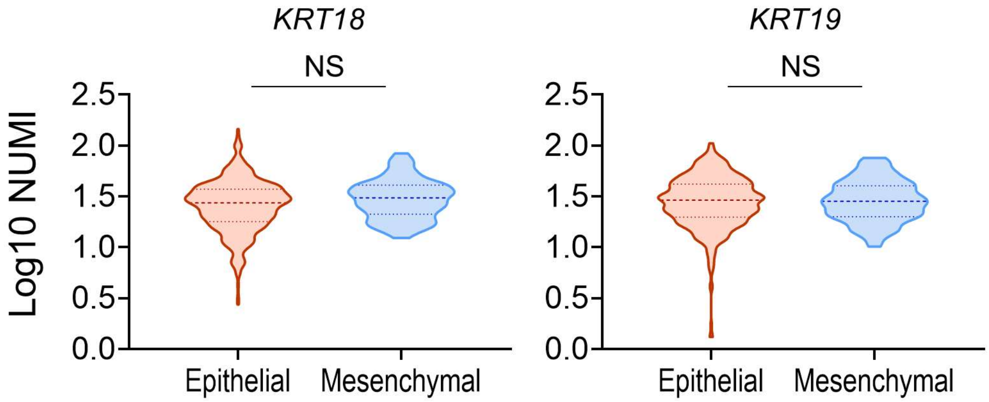


**Figure S7.** Comparison of expression levels of *KRT18* and *KRT19* between epithelial and mesenchymal CTCs. The dash and dot lines of each violin plot denote the median and first and third quartiles, respectively (NS: not significant).

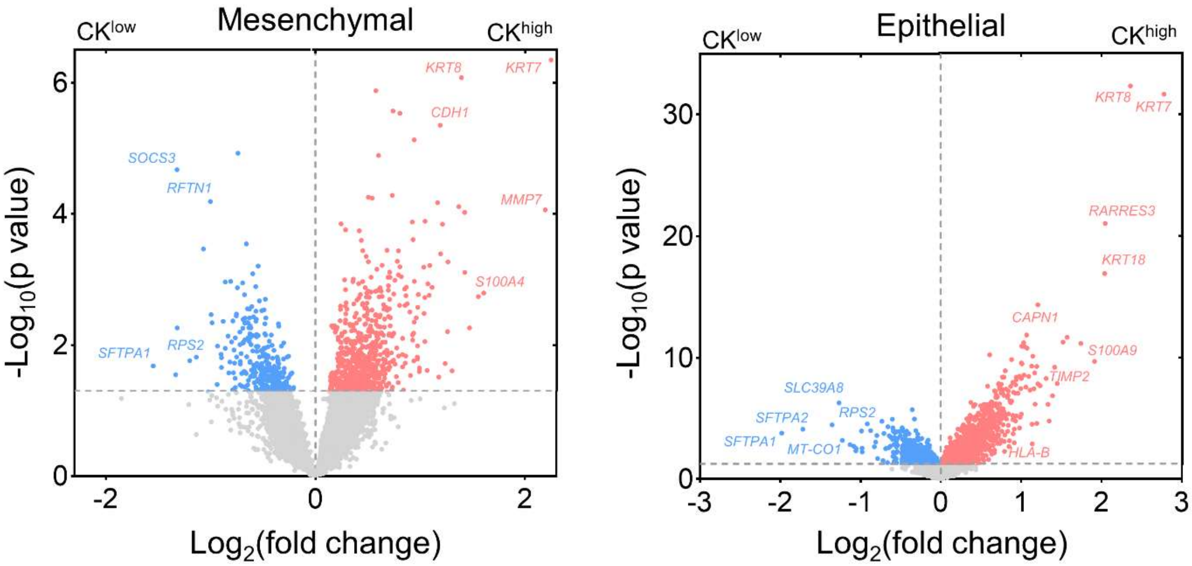


**Figure S8.** Volcano plots showing the DEGs between $CK^{high}$ and $CK^{low}$ CTCs in the epithelial and mesenchymal populations. Statistically insignificant DEGs ($p>0.05$) are colored in grey.

### Supplementary Tables

**Table S1.** Clinical information and CTC counting results of blood samples from treatment-naïve LUAD patients in this study.

| No | Age/Sex | Stage | # of CTC subtypes | | | # of total CTCs | $CK^{neg}$ CTC (%) |
|---|---|---|---|---|---|---|---|
| | | | $HK2^{high}/CK^{neg}$ | $HK2^{low}/CK^{pos}$ | $HK2^{high}/CK^{pos}$ | | |
| 1 | 72/F | IV | 51 | 1 | 18 | 70 | 73 |
| 2 | 68/F | IV | 1 | 117 | 2 | 120 | 1 |
| 3 | 64/M | IV | 10 | 0 | 0 | 10 | 100 |
| 4 | 61/F | IV | 8 | 3 | 0 | 11 | 73 |
| 5 | 72/F | IV | 5 | 0 | 0 | 5 | 100 |
| 6 | 48/F | IV | 3 | 0 | 0 | 3 | 100 |
| 7 | 67/F | IV | 0 | 2 | 0 | 2 | 0 |
| 8 | 50/F | IV | 0 | 0 | 5 | 5 | 0 |
| 9 | 55/M | IV | 7 | 10 | 0 | 17 | 41 |
| 10 | 60/F | IV | 0 | 3 | 0 | 3 | 0 |
| 11 | 70/M | IV | 2 | 0 | 0 | 2 | 100 |
| 12 | 66/F | IV | 3 | 0 | 0 | 3 | 100 |
| 13 | 38/F | IV | 1 | 0 | 1 | 2 | 50 |
| 14 | 61/F | IV | 0 | 1 | 0 | 1 | 0 |
| 15 | 61/M | IIIB | 3 | 0 | 3 | 6 | 50 |
| 16 | 66/M | IIIA | 8 | 0 | 0 | 8 | 100 |
| 17 | 63/F | IV | 0 | 0 | 0 | 0 | - |
| 18 | 54/F | IV | 0 | 0 | 0 | 0 | - |
| 19 | 54/F | IV | 0 | 0 | 0 | 0 | - |
| 20 | 67/F | IV | 0 | 0 | 0 | 0 | - |
| 21 | 68M | IV | 0 | 0 | 0 | 0 | - |
| 22 | 56/M | IV | 0 | 0 | 0 | 0 | - |
| 23 | 49/F | IV | 0 | 0 | 0 | 0 | - |
| 24 | 68/M | IIIB | 0 | 0 | 0 | 0 | - |

**Table S2.** Clinical information and CTC counting results of MPE samples from treatment-naïve LUAD patients in this study.

| No | Sample type | Age/Sex | Stage | Volume (mL) | # of CTC subtypes | | $CK^{neg}$ CTC (%) |
|---|---|---|---|---|---|---|---|
| | | | | | $HK2^{high}/CK^{neg}$ | $HK2^{high}/CK^{pos}$ | |
| 25 | MPE | 70/M | IV | 5 | 25 | 100 | 20 |
| 26 | MPE | 75/M | IV | 10 | 12 | 355 | 3.3 |
| 27 | MPE | 84/F | IV | 5 | 32 | 3939 | 0.8 |
| 28 | MPE | 73/M | IV | 0.7 | 18 | 11760 | 0.2 |
| 29 | MPE | 67/F | IV | 5 | 32 | 3040 | 1.1 |
| 30 | MPE | 65/F | IV | 1.5 | 9 | 260 | 3.5 |

**Table S3.** Clinical information and CTC counting results of CSF samples from treatment-naïve LUAD patients in this study.

| No | Sample type | Age/Sex | Stage | Volume (mL) | # of CTC subtypes | | | $CK^{neg}$ CTC (%) |
|---|---|---|---|---|---|---|---|---|
| | | | | | $HK2^{high}/CK^{neg}$ | $HK2^{high}/CK^{pos}$ | $HK2^{low}/CK^{pos}$ | |
| 31 | CSF | 52/M | IV | 1.5 | 8 | 258 | 1564 | 0.44 |
| 32 | CSF | 37/F | IV | 1.5 | 11 | 202 | 77 | 3.8 |
| 33 | CSF | 51/F | IV | 1.5 | 3 | 11 | 114 | 2.3 |

**Table S4**. Primers used in this study. All primers were synthesized by Genewiz.

| Primer Name | Sequence (5'-3') |
|---|---|
| EGFR-exon19-F | GTGGCACCATCTCACAATT |
| EGFR-exon19-R | ATGCTCCAGGCTCACCAAG |
| EGFR-exon21-F | TTCGCCAGCCATAAGTCCT |
| EGFR-exon21-R | TCATTCACTGTCCCAGCAAG |
| Chr1F | TTTAGGCGTCATCTGAGGGTA |
| Chr1R | TGGCAGCAGTATGGAGAATGTA |
| Chr2F | AGCGGGAGGGACTATTTCAC |
| Chr2R | GGATCGTTCAAAGGGAACTG |
| Chr3F | CCCTTGTACTGGCTCGTGTT |
| Chr3R | CTTGCACATGAAGGTCTGGA |

| | |
|---|---|
| Chr4F | GAGCATCTCTTGGCTCTGCT |
| Chr4R | TTGGGAAAGCACAGATCCTT |
| Chr5F | ACGGACAGTGGACAGATTGC |
| Chr5R | CCACTGTGCCACCCCATT |
| Chr6F | GAGGAGGGCAAGGAGAGAGT |
| Chr6R | ACCCTCCAGTGTGCAAAAAC |
| Chr7F | CTTCCTGCCATTCCACAAGT |
| Chr7R | CCCACTTTCATGCCTCTGAT |
| Chr8F | CTTCCCTGCCTTGCTCTCTA |
| Chr8R | CGGGACATTTCAGCAATCTT |
| Chr9F | CTGTGGAGCAGCTGTTTCTG |
| Chr9R | GAATTCACAAAGCCCCAAGA |
| Chr10F | CCCCTCATTCAAATCAGCAT |
| Chr10R | CAGGCAAAAGCTGGAGTTTC |
| Chr11F | TGAATGAGAACGCAGATGTGA |
| Chr11R | CACAAAGCATCCAGGGTCATT |
| Chr12F | ATCATGGAAATGCAGCCTCT |
| Chr12R | AGAACCCAGCTGGAATGATG |
| Chr13F | TGTTTCATGGAGTCCTGCTG |
| Chr13R | GGAGGCAAGAACCAAACAAA |
| Chr14F | AGCCAAGACGTACCCTCTCA |
| Chr14R | TGCTTTACACCAATCCCACA |
| Chr15F | TCAGCATGGGTTATGGGTTT |
| Chr15R | CCCAGATGATGGAGAGGAAA |
| Chr16F | GCCTGTGTTTGCTGATGAAA |
| Chr16R | GGGCAACGACCGTACTTAAA |
| Chr17F | TCCTGGGCTAGCCTTTTACA |
| Chr17R | ATCGCTTGAGCACTGAAGGT |
| Chr18F | AGACGAGCCTTTCTCTGTCG |
| Chr18R | TCGAGACCATCCCCACTAAC |
| Chr19F | AGTTGAGGAGATGGTGGAGC |
| Chr19R | AACAGGAGCCTTGGTCAGTC |
| Chr20F | CTGGTCAAACATCTCCCTCGT |
| Chr20R | CTCCACGCATCTTACATCACCT |
| Chr21F | GGACTTTGCTGACGGGATTA |
| Chr21R | GAACTAACGACCTCACGCTTG |
| Chr22F | TCCACAACCCCTTATCTTACCC |
| Chr22R | ACCTCAGGTGATCTACCCGC |

**Supplementary Datasets**

**Dataset S1**. Expression of the 200 EMT-defining genes and EMT enrichment scores of all the CTCs and TCGA patient samples.

**Dataset S2.** List of top 100 DEGs upregulated in $CK^{high}$ (or $CK^{low}$) cells in epithelial and mesenchymal CTC populations.
